# Principles of transcription factor traffic on folded chromatin

**DOI:** 10.1101/164541

**Authors:** Ruggero Cortini, Guillaume Filion

## Abstract

All organisms regulate the transcription of their genes. To understand this process, it is essential to know how transcription factors find their targets in the genome. In addition to the DNA sequence, several variables have a known influence, but overall the binding patterns of transcription factors distribution remains mostly unexplained in animal genomes. Here we investigate the role of the chromosome conformation in shaping the search path of transcription factors. Using molecular dynamics simulations, we uncover the main principles of their diffusion on folded chromatin. Chromosome contacts play a conflicting role: at low density they enhance the traffic of transcription factors, but a high density they lower the traffic by volume exclusion. Consistently, we observe that in human cells, highly occupied targets, where protein binding is promiscuous, are found at sites engaged in chromosome loops within uncompact chromatin. In summary, those results provide a theoretical framework to understand the search trajectories of transcription factors and highlight the key contribution of genome conformation.

## 1 Introduction

Transcription factors play a key role in the regulation of transcription [Levine and Tjian, 2003]. Upon binding cognate regulatory sequences, they trigger a cascade of molecular events leading to the recruitment of the RNA polymerase at promoters and its maturation into a processive elongation complex [Saunders et al., 2006]. Understanding how transcription factors find their targets in the genome is thus the very first step to understanding how transcription is controlled at the molecular level.

Transcription factors contain a DNA binding domain that usually has a non-specific affinity for DNA and a specific (high) affinity for some target sequence called the binding motif [Stormo, 2000]. Affinity for their motifs predicts the binding of transcription factors *in vitro* [Badis et al., 2009], but the ChIP-on-chip and ChIP-seq technologies have painted a different picture in the nucleus of multicellular eukaryotes. Transcription factors occupy sites where their motif is absent [Moorman et al., 2006, Kvon et al., 2012], and leave unbound most sites where it is present [Farnham, 2009, Schmidt et al., 2010, Kaplan et al., 2011, Arvey et al., 2012]. This indicates that a simple protein-DNA interaction model is insufficient to describe transcription factor dynamics *in vivo*.

The nucleus is a rich and hetereogenous environment, where higher order interactions between molecules take place. For instance, the presence of histones on most of the genome is a barrier to the binding of transcription factors. Related to this constraint, the so called “pioneer transcription factors” can bind their target in the presence of a nucleosome and help other transcription factors access their targets [Zaret and Carroll, 2011]. However, the binding of pioneer transcription factors is vastly different between cell types [Hurtado et al., 2011], conflicting with the simple view that they bind their motif regardless of the chromatin context. Additional phenomena dictate the search process of transcription factors in the nucleus.

Among them is a mechanism known as “facilitated diffusion” [Berg et al., 1981, von Hippel and Berg, 1989], which emphasizes the key role of non-specific affinity for the search kinetics. Transcription factors are first adsorbed onto DNA or chromatin through an electrostatic pull acting at short distance [Dahirel et al., 2009], and for a short period of time they diffuse on the polymer. This “scanning” or “sliding” mode is essential to discover the target sequence [Berg et al., 1981, Mirny et al., 2009]. However, both terms are potentially misleading: transcription factors may merely detach and reattach to the chromatin immediately. At the points where chromatin fibers meet, transcription factors can thus fall off one fiber and reattach to the other, effectively jumping over large genomic distances while diffusing on chromatin. The three-dimensional conformation of naked DNA is known to affect the search kinetics *in vitro* [van den Broek et al., 2008], but little is known about similar search mechanisms in chromatin.

The importance of this mechanism of diffusion is well established in the eukaryotic nucleus [Tafvizi et al., 2008] and it is now clear that transcription factors diffuse intermittently in the nucleoplasm and on the chromatin fiber [Izeddin et al., 2014, Normanno et al., 2015]. However, little is known about the impact of the geometry of the chromatin fiber. Genomes have a characterstic three-dimensional structure, revealed by chromosome capture-based methods such as Hi-C [Lieberman-Aiden et al., 2009]. It is still unclear how genomes acquire a particular conformation, but once preferential contacts are established, they can influence the trajectories of transcription factors diffusing on chromatin.

It is thus tempting to think that the knowledge of chromosome conformation can be used to obtain some insight into the dynamics of transciption factor search. However, the general principles are presently unknown for lack of a theory. Previous work by ourselves and others suggested methods to infer the binding profiles of proteins on DNA from Hi-C matrices [Corrales et al., 2017, Avcu and Molina, 2016, Wang et al., 2017], but the validity of those approaches remains unproven because there is no guarantee that they correspond to realistic physical processes.

Here we establish the basic principles of transcription factor diffusion on folded chromatin. We use a molecular dynamics simulation approach to investigate the role of chromosome conformation in the search process. Exploring configurations with the strings and binders model of Barbieri et al. [2012], we find that the geometry has a large influence on the traffic of diffusing bodies with an affinity for the polymer. Strikingly, polymer loops increase traffic and occupancy in a wide range of conditions, but decrease them at high compaction. Consistently, Hi-C and ChIP-seq data show that massive protein binding accumulates at genomic loci engaged in long distance contacts. Overall, this work suggests that the three-dimensional conformation of the genome affects the discoverability of binding sites and contributes to the global distribution of transcription factors.

## 2 Results

We used molecular dynamics simulations to understand how the conformation of the genome can affect the diffusion of transcription factors. This modeling strategy captures the behavior of simple objects evolving according to realistic physical interactions. To develop a general and tractable model we abstracted the specific features of chromatin and transcription factors to their bare essentials: Chromatin was considered as a folded polymer and transcription factors as diffusible molecules with an affinity for chromatin.

Transcription factors were represented by spherical particles referred to as *tracers*. Most importantly, tracers differ from transcription factors by the fact that they do not have targets, *i.e.* they have no specific affinity for a particular region of the polymer. They only interact with the polymer through uniform non-specific affinity. This aspect of the model is essential to capture the dynamics of transcription factors in search mode, rather than in bound mode. In other words tracers can be thought of as non-specific transcription factors, or as transcription factors not bound to their target site. We labelled *ε* the strength of the non-specific affinity of the tracers for the polymer.

The folded polymers were simulated using a model originally developed by Barbieri et al. [2012] and summarized graphically in Figure 1. Briefly, the model describes the large-scale structure of the polymer as an aggregate of stable loops formed between predefined anchor monomers. The loops are formed by special particles called *binders* that have a high affinity for anchor monomers and can thus bridge them together. The overall openness or compaction of the polymer depends on the number of loops, or more accurately on the fraction of these anchor monomers, labelled *ϕ*.

**Figure 1:**
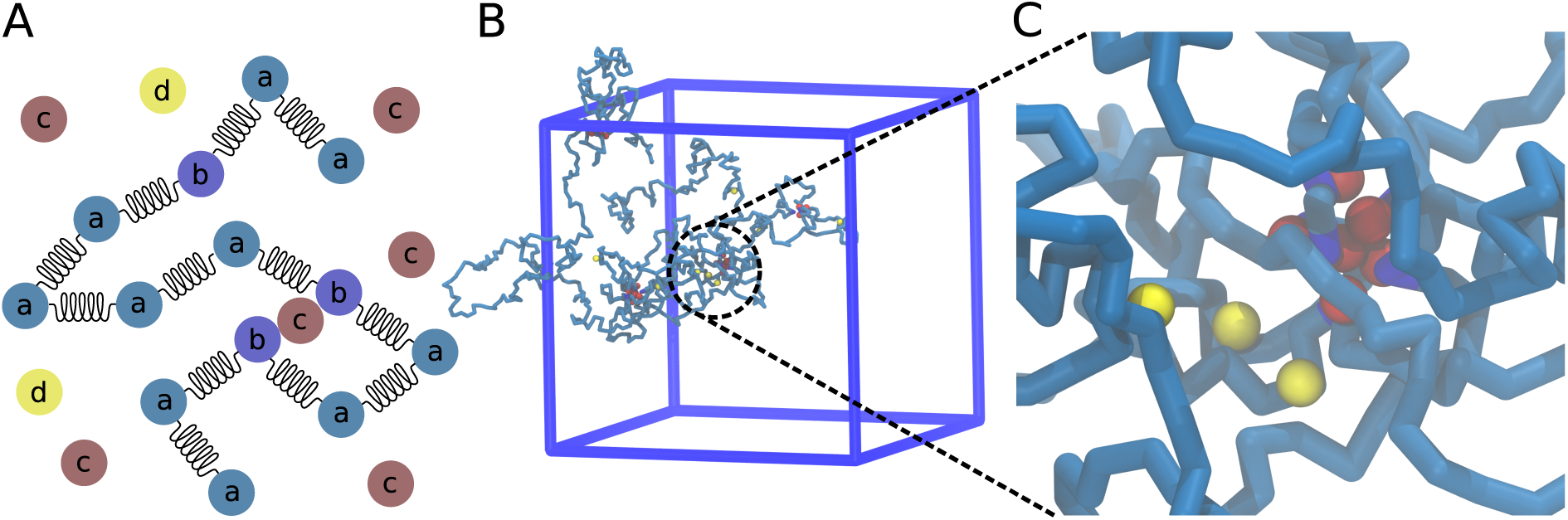
Description of the simulation setup. (A) The polymer is made of a and b particles, connected by harmonic springs. Binders (c particles) are introduced to bridge b particles. Tracers (d particles) interact non-specifically with both a and b particles. Binders and tracers interact only by hardcore repulsion, *i.e.* by volume exclusion. (B) snapshot of a simulation together with the simulation box. (C) zoom in of a loop formed by the binders and their binding sites, along with the yellow tracers bound nearby.

Varying *ε* and *ϕ* allowed us to explore a wide range of conditions. For each simulation, we computed the intra-polymer contact matrix where each entry was the number of times two given monomers were in contact during the simulation (meaning that their distance is less than a threshold *t*, see Methods). From this matrix, we computed the row sum, denoted ***R***, that represents the total amount of contacts for each monomer. We also computed the total number of contacts between the tracers and the monomers, denoted ***C*** and referred to as the occupancy profile of the tracers. This profile represents the total amount of time tracers spent in contact with different regions of the polymer.

We carried out two independent sets of simulations, one with 10 tracers and another one with 200 tracers. This allowed us to test the effect of the tracers themselves on the conformation of the polymer.

### 2.1 Polymer loops have two opposite effects on the traffic of the tracers

Figure 2 illustrates the behaviour of the system when the parameters take their extreme values. The main point is to notice the correspondence between the monomer contact frequency (array ***R***) and the occupancy profile of the tracers (array ***C***). The correspondence is particularly clear in the cases with only a few loops in the polymer (low *ϕ*) and strong binding of the tracers (high *ε*).

**Figure 2:**
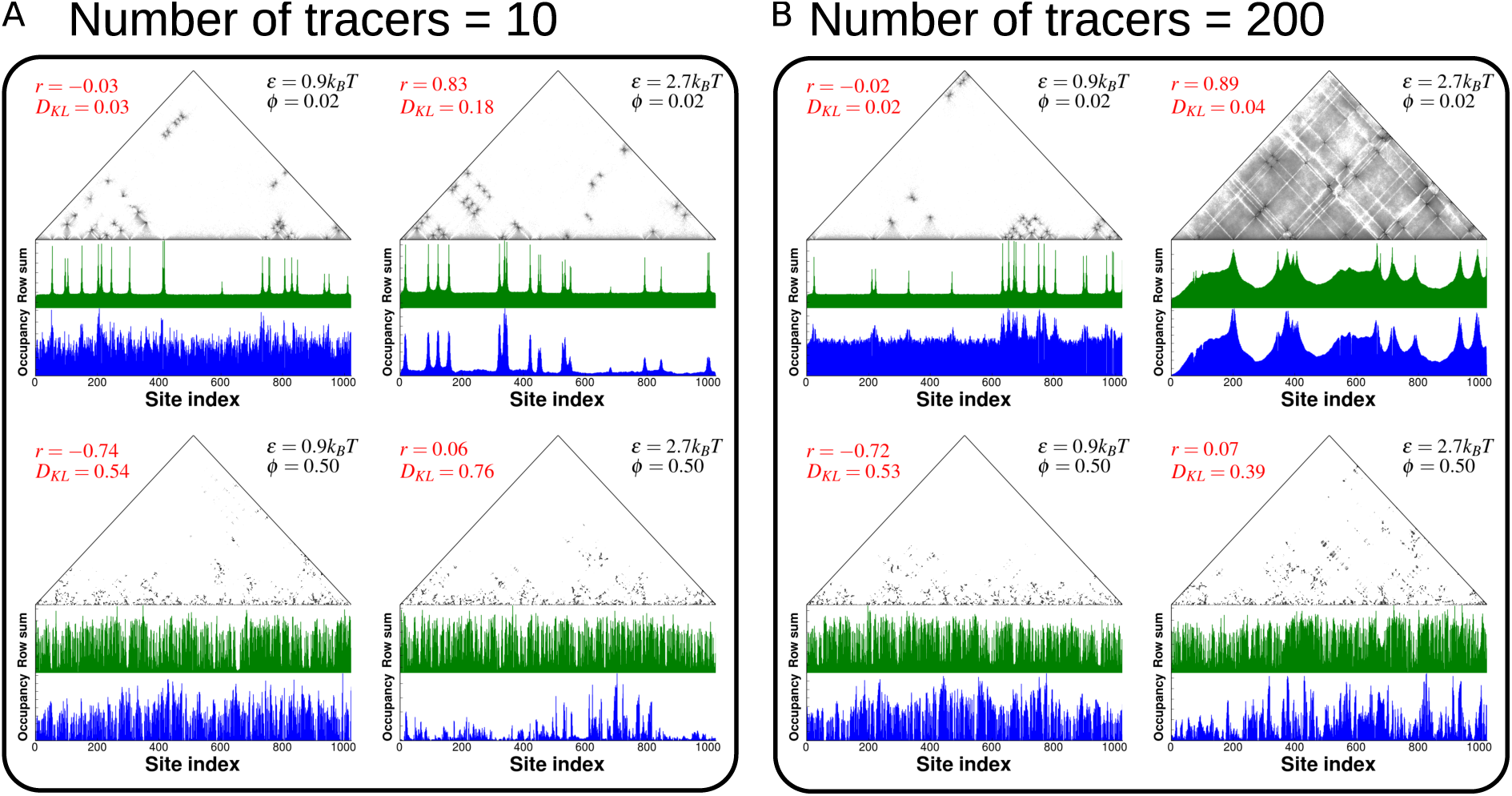
Comparison of single simulation results for the cases of *n*_*t*_ = 10 (panel A) and *n*_*t*_ = 200 (panel B). The examples refer to the smallest and largest values of *ϕ* and *ε* in all their combinations. For each example, we show the contact matrix of the polymer (in log scale), the occupancy profile of the tracers (***C***, “Occupancy”) and ithe total amount of contacts for each monoer (***R***, “Row sum”). Each plot indicates the values of the Pearson correlation coefficient *r* between ***C*** and ***R***, along with its Kullback-Leibler divergence *D*_*KL*_(***C****|****R***).

We measured the correspondence between ***C*** and ***R*** with the Pearson correlation coefficient *r* and the Kullback-Leibler (KL) divergence *D*_*KL*_(***C*** *|* ***R***). These two measures are both valid metrics but they represent fundamentally different features of the data. When using the Pearson correlation coefficient, ***C*** and ***R*** are interpreted as numeric variables and the quantity of interest is their covariation. When using the KL divergence, ***C*** and ***R*** are interpreted as distributions and the quantity of interest is the information lost by using ***R*** instead of ***C***.

In figure 2A we illustrate the behavior of the system with 10 tracers. In this case, the tracers do not affect the conformation of the polymer, as confirmed by measuring its radius of gyration (see Appendix A). If the polymer has few loops (*ϕ* = 0.02) the high values of *r* and the low values of *D*_*KL*_(***C*** *|****R***) indicate that the total amount of contacts coincides with the occupancy profile of the tracers. This is the combination of two effects. First, the *on*-rate of the association reaction is doubled at the contact points because they come from two distinct branches of the polymer. Second, the *off* -rate is reduced because the local concentration of the polymer increases, thereby favouring binding (see Supplementary Movie 1).

Highly compacted polymers with a high number of looping sites (*ϕ* = 0.50), define another regime. High compaction combined with low tracer-polymer affinity (low *ε*) result in a strong *anti*correlation between the amount of contacts and the occupancy profile of the tracers, *i.e.* the tracers tend to visit the sites that make *fewer* contacts. The reason is that the polymer forms a globule and the most visited monomers are at the surface, where their contacts with other monomers are less frequent. In comparison, when the tracer-polymer affinity is high (high *ε*) we observe a “caging” effect, whereby the tracers remain blocked in compacted inner structures inside the polymer (see below). In this case both *r* and *D*_*KL*_(***C R***) report that ***C*** and ***R*** are unrelated (see Supplementary Movie 2).

Figure 2B shows the results for 200 tracers. At low compaction (low *ϕ*) the polymer undergoes a transition from coil-like to globular as the tracer-polymer affinity increases (see Appendix A). This means that the tracers are themselves creating the contacts between distal polymer segments. Consistently, the occupancy profile of the tracers is in close agreement with the number of contacts of the monomers. The comparison with panel A immediately shows that many additional intra-polymer contacts are induced by the tracers themselves. When *ϕ* is high, the polymer is already globular, so the shape does not change as *ε* increases. The results are otherwise similar to the simulations with 10 tracers.

Figure 3 summarizes these results with the average values of the KL divergence and the Pearson correlation coefficient for all the parameters tested in our simulations. Regardless the number of tracers, a larger number of loops results in a poorer correspondence between ***C*** and ***R***. In summary, the conformation of the polymer defines two regimes. In the first, the polymer is uncompact and contacts predict the traffic of the tracers, in the second, the polymer is compact and contacts do not predict the traffic of the tracers.

**Figure 3:**
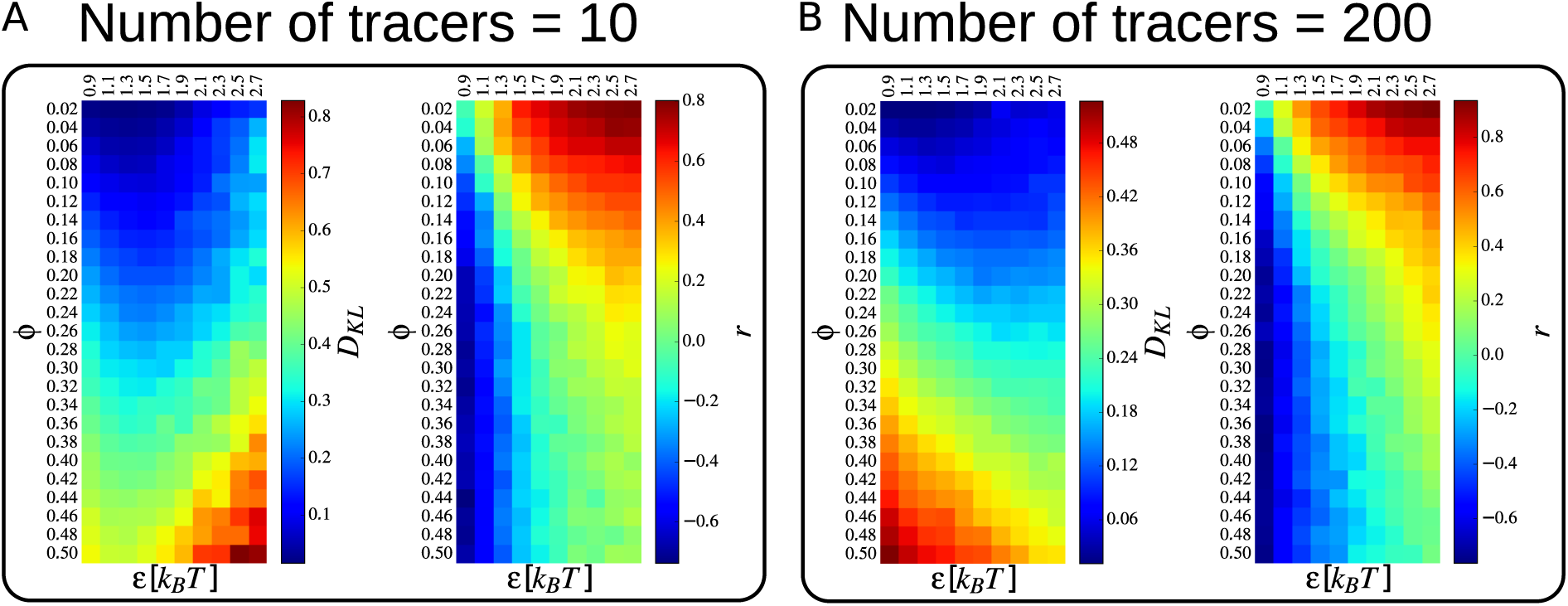
Average values of the Kullback-Leibler divergence and of the Pearson correlation coefficient when comparing the row sum of the polymer contact matrix and the occupancy of the tracers (see also figure 2). (A) in the case of *n*_*t*_ = 10; (B) *n*_*t*_ = 200. All calculations were performed as described in Methods.

### 2.2 High polymer compaction excludes tracers

Why does the conformation of the polymer cease to predict the traffic of the tracers as the number of loops increase? At least two non mutually exclusive scenarios can be imagined. In the first, the polymer forms globular domains that are too dense for tracers to enter. In the second, tracers enter the domains but steric effects prevent them from binding the anchors of the loops.

We defined the coverage of the polymer as the percentage of monomers visited by a tracer at least once during the simulation. Figure 4 shows that the coverage decreases as the number of loops increases (high *ϕ*), consistent with the view that the tracers cannot access the core of the polymer. However, observe that increasing the affinity of the tracers alleviates this effect, so the exlusion does not proceed by hardcore repulsion. In fact most of the monomers are accessible at some value of the non-specific affinity, so the polymer is never so dense as to be completely impermeable to tracers.

**Figure 4:**
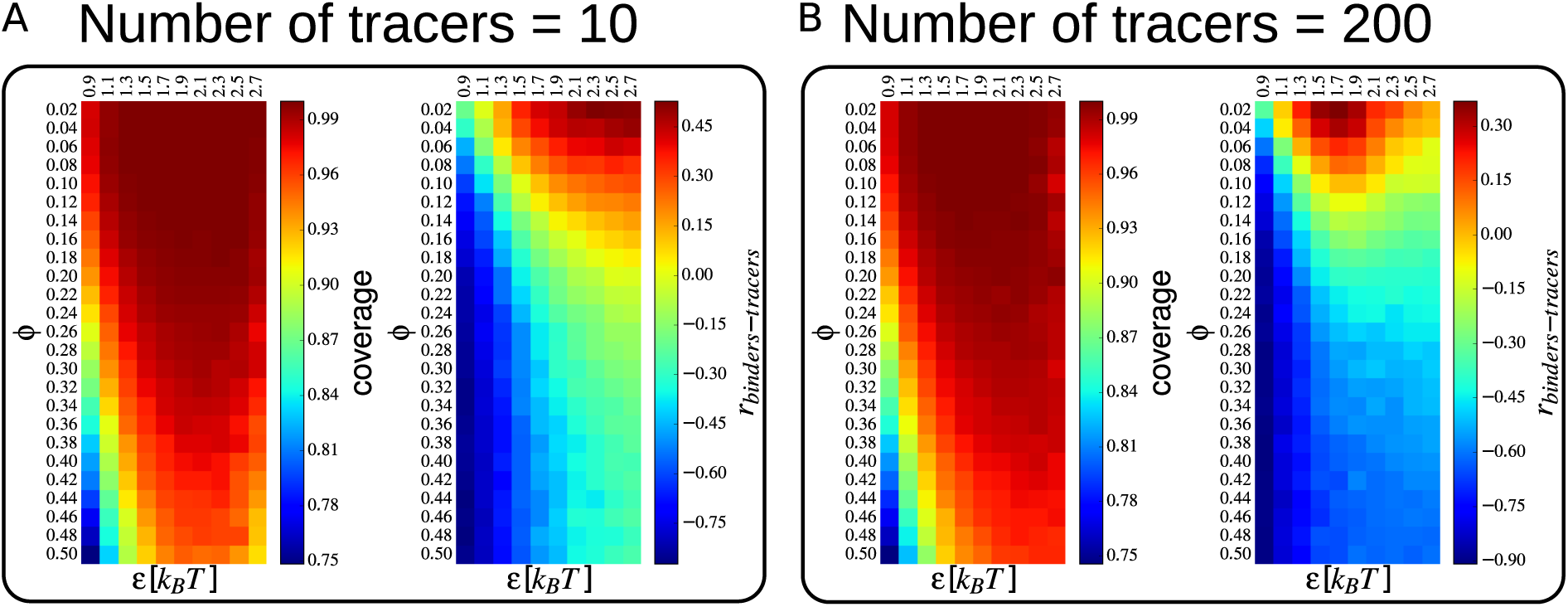
Volume exclusion effects in the simulations. For the two simulation sets with *n*_*t*_ = 10 (A) and *n*_*t*_ = 200 (B), we plot the average values of the Pearson correlation between the tracer and binder occupancy sets, and the average percentage of sites of the polymer that are visited by the tracers during the simulations (coverage).

We also computed the average correlation between the occupancy of the tracers and the occupancy of the binders. Recall that the binders bridge the loop anchors and remain fixed on the same monomers for the duration of the simulation. As the number of loops increases (high *ϕ*) the correlation of the occupancy profiles becomes strongly negative, indicating that the binders exclude the tracers at the looping sites. Once again, part of the effect can be alleviated by a higher non-specific affinity, but the correlation remains negative at high number of loops.

These results suggest that both effects apply upon polymer compaction. The core of the polymer domains becomes less accessible and the anchor sites become crowded. As a result, contacts within the polymer become poor predictors of the occupancy of the tracers. These results were also confirmed by calculating the contact enrichment between the tracers and the anchor sites (see Methods and figure 10).

We confirmed the effect of volume exclusion by simulating tracers without affinity for the polymer (only hard core repulsion). In these conditions, the equilibration time is prohibitive with molecular dynamics simulations, so we turned to Monte Carlo simulations. With this approach, we could test the importance of the attractive tracer-polymer interactions in determining the occupancy patterns (see Methods).

Figure 5 shows that we obtained opposite results at low and high compactions. When *ϕ* = 0.02 (low compaction), the results from the molecular dynamics simulations (performed at *ε* = 0.9*k*_*B*_*T*) and from the Monte Carlo simulations are mostly unrelated (*r* = 0.04). This confirms that the attractive interactions between the tracers and the polymer are crucial in determining the occupancy profile.

**Figure 5: figure.**
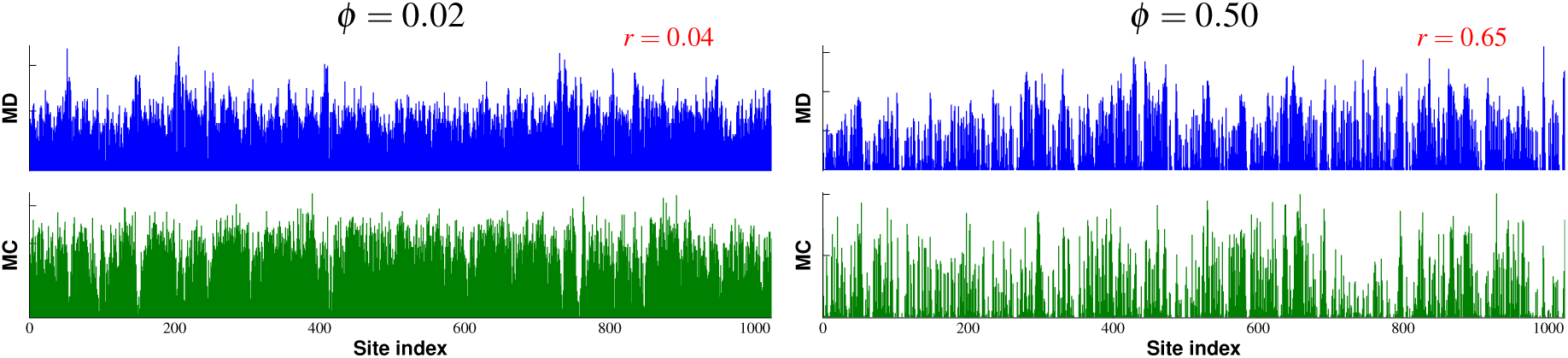
Occupancy obtained by Molecular Dynamics (MD) or Monte Carlo (MC) simulations. In molecular dynamics simulations, the affinity of the tracer is equal to 0.9*k*_*B*_*T* and in Monte Carlo simulations, the tracters have no affinity for the polyer (they interact with it only through hard core repulsion). Left: Low compaction with *ϕ* = 0.02. Right: High compaction with *ϕ* = 0.50.

When *ϕ* = 0.50 (high compaction), the occupancy profiles of tracers with or without affinity for the polymer are similar (*r* = 0.65). This shows that in this regime, the affinity between the polymer and the tracers has little impact on the occupancy.

### 2.3 Highly occupied targets in the human genome

An important question is whether the phenomena observed in our simulations are relevant to living cells. There is general agreement that chromatin compaction tends to exclude transcription factors [Bancaud et al., 2009], but the idea that the geometry of chromatin may guide transcription factors is still exploratory [Benichou et al., 2011, Brackley et al., 2012, Avcu and Molina, 2016, Wang et al., 2017]. Our results indicate that at low chromatin compaction, long-range chromosomal contacts may increase the traffic of transcription factors. The simulations predict a 2-5 fold increase of non-specific binding (figure 2). The effect is small, but it applies to every molecule with an affinity for chromatin.

We conjectured that loop-enhanced traffic may explain one of the least understood patterns of transcription factor occupancy, namely the highly occupied targets [HOTs, Moorman et al., 2006, Kvon et al., 2012, Foley and Sidow, 2013]. HOTs or binding hotspots are small regions of the genome (typically less than 1 kb) bound by most proteins with an affinity for chromatin. The promiscuous binding of transcription factors in HOTs is independent of the presence of their binding motifs, suggesting that the process depends on non-specific affinity and protein-protein interactions. Could HOTs result from the formation of chromatin loops?

To test this hypothesis, we used high-resolution Hi-C data performed in the human cell line GM12878 [Rao et al., 2014], together with the locations of HOTs obtained from the ChIP-seq profiles of 96 proteins mapped in the same cell line [Foley and Sidow, 2013]. Visually, HOTs tend to localize at regions engaged in long distance chromosomal contacts (figure 6A). More generally, HOTs are strongly enriched at loop anchors, as defined by Rao et al. [2014]: 2,138 of the 12,887 HOTs lie at the basis of a loop, compared to 528 expected (figure 6B). Thus the trend is very robust, especially considering the imprecision in calling loops and HOTs.

**Figure 6:**
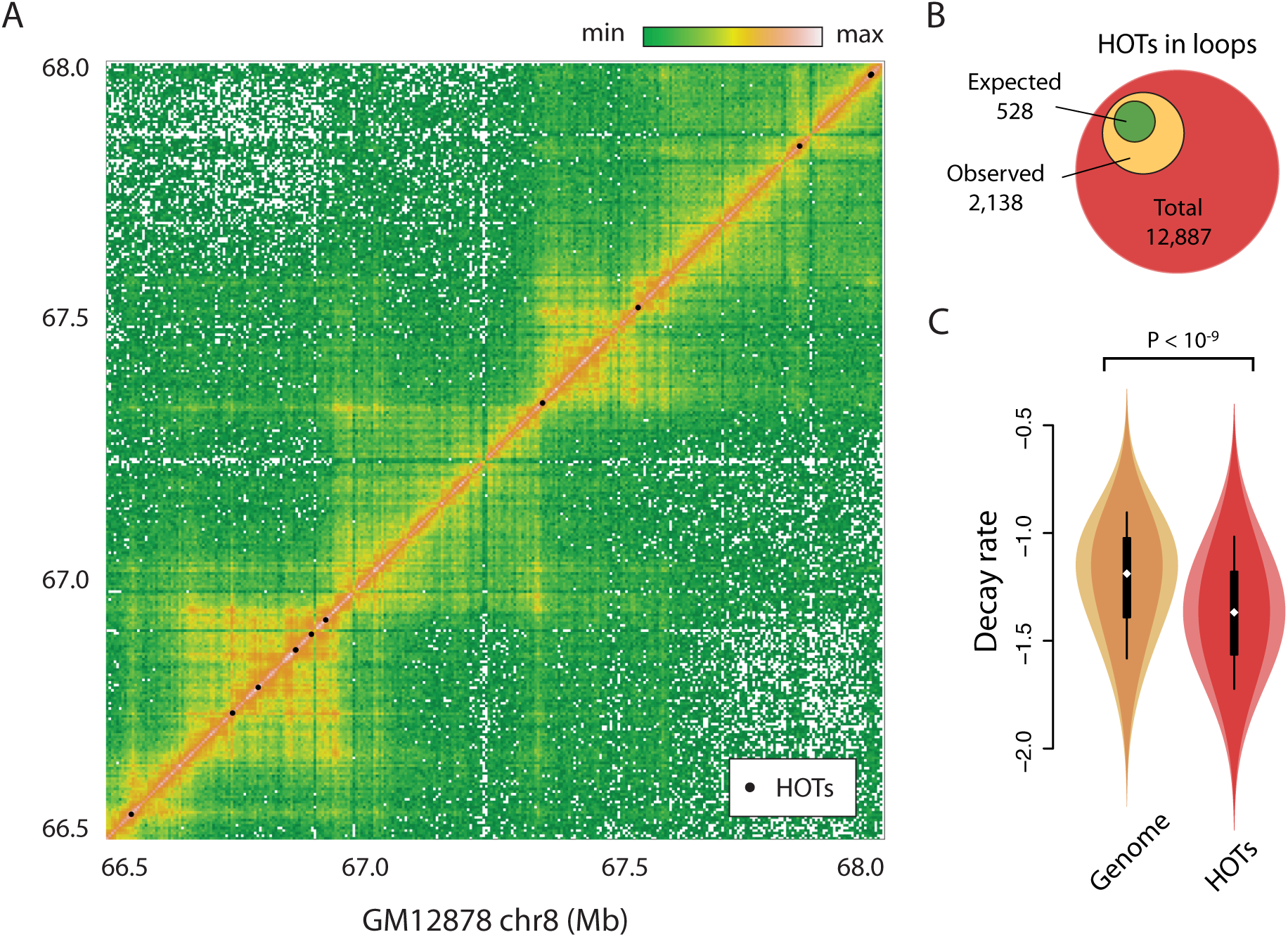
Highly occupied targets in the human genome. (A) Hi-C map of the human cell line GM12878, along with) the position of the highly occupied targets (HOTs) indicated by black dots. (B) HOTs are enriched at chromosomal loops in GM12878. The location of the HOTs from Foley and Sidow [2013] was crossed with the location of the loops from Rao et al. [2014]. Out of 12,887 HOTs, 2,138 were found in loops versus 528 expected (binomial distribution *P <* 10^-9^). (C) HOTs are located in uncompact chromatin. The violin plots show the distribution of contact decay in 50 kb windows in the whole genome versus each HOT. The mean values are -1.22 and -1.37, respectively (95% confidence interval for the difference 0.146-0.157, Student’s *t* test with Welch-Satterthwaite correction and 18,080 degress of freedom *P <* 10^-9^).

The simulation results also predic that HOTs should be present in the regions of the genome that are the least compact. The standard way to compare compaction levels is to estimate the local rate of contact decay, which is a measure of the polymer state [Barbieri et al., 2012]. Hi-C contacts are locally proportional to *s* ^*α*^, where *s* is the linear separation between the loci and *α* is the decay rate. Values of *α* that are close to 0 indicate compact regions, whereas strongly negative values indicate open regions. The local contact decay around HOTs in the data set is significantly lower than the genome average (figure 6C). This indicates that HOTs tend to occur in the least compact regions of the genome, consistent with the simulation results.

Overall, these results indicate that massive protein binding tends to accumulate at loop anchors in open regions of the nucleus, consistently with our prediction that chromatin loops can enhance the traffic of transcription factors.

## 3 Discussion

In this study we used molecular dynamics simulations to establish the governing principles of transcription factor diffusion on folded chromatin (figure 7). The main feature of the polymer structure is the amount of contacts, which acts in two opposite ways (figure 3): At low density, it increases the traffic of the tracers; at high density, the compaction of the polymer induces volume exclusion effects that keep the tracers away from loop anchors (figure 4). In the latter case, the tracers tend to interact with the monomers on the outside of globular domains, but they can also enter the structure, especially if their affinity for the polymer is high. These results are in line with experimental data obtained in live cells [Bancaud et al., 2009], where it was shown that chromatin compaction excludes diffusible factors, but that no compartment remains fully inaccessible.

**Figure 7:**
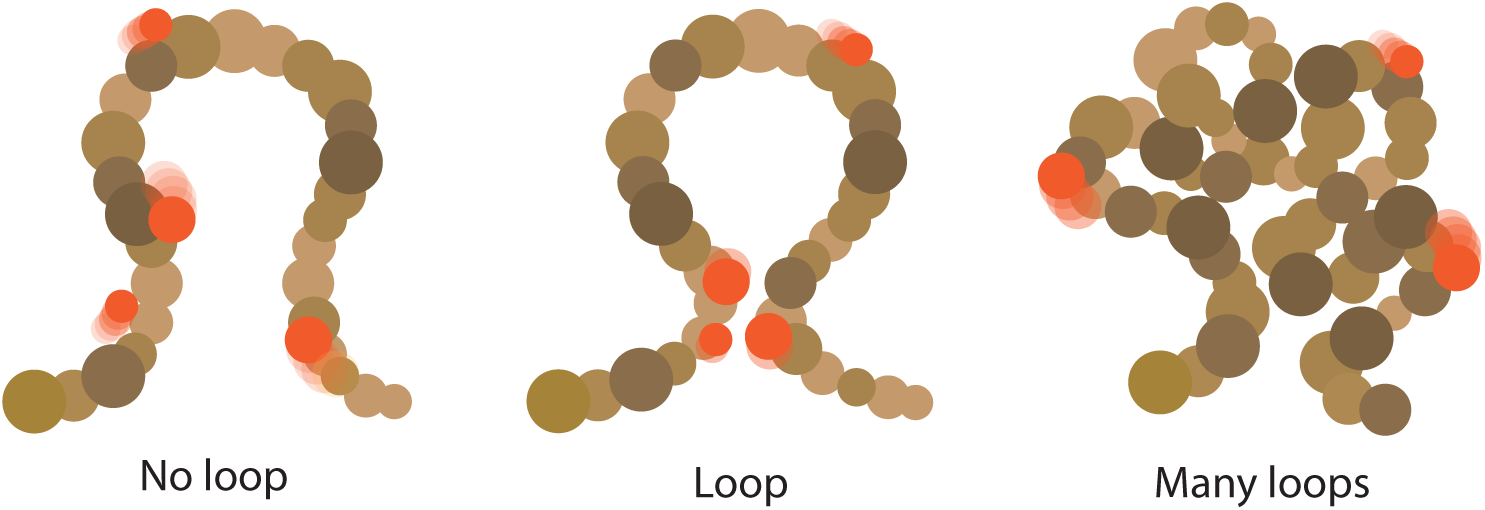
Effect of chromosome conformation on the traffic of transcription factors. At low compaction (left), transcription factors slide uniformly on the chromatin fiber. In the presence of a loop (middle), transcription factors accumulate at the contact point. When the number of loops is high (right), transcription factors diffuse on the outer shell of the globule due to volume exclusion effects.

Importantly, these effects describe the behavior of transcription factors in search mode only. Chromatin loops may increase the *on* rate by a factor 2-5 (the number of fibers at the contact point, see figure 2), but the *off* rate may vary by several orders of magnitude with the DNA sequence [Badis et al., 2009]. A 2-fold effect is large for the binding kinetics, but negligible for the average occupancy of high affinity binding sites. Consistently, it was shown that the target sites of many transcription factors remain bound on the mitotic chromosome [Teves et al., 2016], where the conformation is radically different [Naumova et al., 2013].

Nevertheless, transcription factors spend a large fraction of the time in search mode [Izeddin et al., 2014, Normanno et al., 2015] and they accumulate at highly occupied targets (HOTs), where most binding motifs are not found [Kvon et al., 2012]. Our results suggest that this alternate binding mode depends on the conformation of the genome. The HOTs are found primarily at loop anchors in uncompact regions (figure 6), as predicted by the simulations (figures 2 and 3). The key point is that chromosome conformation effects apply to all the molecules in the nucleus with an affinity for chromatin. This permits the appearance of emergent behaviors, induced by the interactions between the components.

### 3.1 Transcription factors create the contacts

In the molecular dynamics simulations, the structure of the polymer is “given” by the fixed distribution of the binders. This model recapitulates the essential features of Hi-C maps [Barbieri et al., 2012], but it does not necessarily correspond to the actual mechanism by which the genome is folded. Actually, this question is still debated [Cortini et al., 2016]. For example, the “loop extrusion” model posits the existence of molecular machines capable of creating loops, which could be cohesins [Fudenberg et al., 2016, Sanborn et al., 2015]. Other authors remarked the crucial role played by epigenetic domains in shaping the three-dimensional genome [Jost et al., 2014].

We showed that when the number of tracers exceed a critical number and a critical value of the affinity, the tracers themselves create intra-polymer contacts. This effect was predicted by the strings and binders switch model of Barbieri et al. [2012], but here we show that it is also valid when the polymer has specific loops in its structure. This indicates that a high concentration of transcription factors may by itself induce the chromatin fiber to collapse locally, a result which suggests that the many transcription factors present in HOTs may stabilize long-range contacts. More work is necessary to clarify the effect of inhomogeneous binding profiles of the tracers/transcription factors in mediating and stabilizing chromatin loops.

A strong assumption that will need to be tested is that transcription factors are close to saturating concentration in the nucleus, so that the weak effect of a chromatin loop is sufficient to nucleate an aggregate. It will also be important to elucidate whether such aggregates are stable or transient, and what other factors are necessary for their formation.

It is known that architectural proteins such as CTCF [Phillips and Corces, 2009] play a crucial role in shaping the three-dimensional genome. The architectural proteins could play the role of binders in our simulations, suggesting that sites in the vicinity of loop anchors have some functional relevance. More studies are needed to unveil these effects.

### 3.2 Genome structure as a guide for complex assembly

It was previously observed that cohesin binding is frequently associated with the presence of many transcription factors [Yan et al., 2013], but the mechanism remained unclear. Since cohesin is necessary for the formation of the loops [Rao et al., 2017], our work suggests the following model: Cohesin binding induces the formation of a chromatin loop, which increases the local concentration of transcription factors. This nucleates the formation of an aggregate stabilized by protein-protein interactions. Finally, the high concentration of transcription factors compacts the chromatin the same way we observed for a large number of tracers, which further stabilize the loop.

The main feature of this interpretation is that it makes the formation of liquid-like aggregates dependent on the geometry of the chromatin polymer. In this regard, the three-dimensional organization of the genome could be a way to guide proteins of the same complex to the same locations to facilitate their assembly. As a side effect, complexes would exist only at the sites where they are needed because they would not assemble without the help of chromatin loops.

This interpretation also suggests that long distance contacts may not be an accidental property of enhancers, but rather their essential mechanism of action. In this regard, enhancers would enable transcription only if they form a contact with another site, thereby promoting transcription factor binding and assembly of the key components of the transcription machinery.

Our model is consistent with the experimental data regarding the distribution of HOTs in the nucleus. It makes several other predictions that can be experimentally tested. The first is that HOTs should disappear in mitosis. Indeed, the compaction of the chromosome and the disappearance of long distance contacts and the loops [Naumova et al., 2013] are unfavourable conditions for the accumulation of transcription factors. The second prediction is that HOTs should disappear or change location if cohesin is destroyed. In the absence of chromatin loops, nucleating the HOTs should happen at random locations or not happen at all. Finally, our model predicts that binding kinetics are 2-3 times faster when the target lies at the basis of a chromatin loop. Since more trajectories lead to the target site when it lies in a loop, a rough estimate is that the search time is reduced by the number of chromatin fibers that meet at the cross point.

As counterintuitive as it may seem, the affinity between a transcription factor and its target may be increased by the formation of a loop. Affinity is often thought of as the strength of the binding at the target site, but it is the ratio of the *on* and *off* rates. As a consequence, any change in the binding kinetics can potentially change the affinity. In this system, kinetic and energetic phenomena are two sides of the same coin so they have to be considered simultaneously.

In summary, this study shows that the conformation of the genome should be taken into account to understand the distribution of transcription factors in the nucleus of animal cells. More generally, the principles highlighted here pave the way for a general theory of facilitated diffusion of transcription factors on folded chromatin.

## 4 Methods

### 4.1 Simulation setup

Our simulation model is basically the same as the strings and binders switch (SBS) model [Barbieri et al., 2012], and consists of the following elements (see figure 1):

- A bead-spring **polymer**, consisting of *N* total particles, which represent the individual monomers. The polymer is made of two types of particles: particles of type “a”, which have no special property, and type “b”, which are binding sites. The fraction of b particles in the polymer is given by the parameter *ϕ*.
- **Binders**, type “c”, which are free to diffuse in space, have strong attractive interactions with the b particles on the polymer. In our system, we keep the number of binders such that there are always two binders per binding site. Therefore, the number of binders is *n*_*b*_ = 2*ϕN*. The interaction strength is chosen in such way that the binders are ordered [Chiariello et al., 2016].
- **Tracers**, which we label “d”, which diffuse freely in space and have an attractive short-ranged interaction with both a and b type particles. The number of tracers is *n*_*t*_.

The polymer is described as a classical bead-spring system. Each particle in our system has a radius *σ*. We chose *σ* = 15nm independently of the particle identity, and *σ* is set to be the length scale in the simulation units. The choice of the length scale is to match more or less the width of the chromatin fiber (30 nm fiber). Successive beads in the polymer are connected with harmonic springs of stiffness *k* = 330*k*_*B*_*T/σ*^2^, as illustrated in Fig. 1. The interaction between successive monomers is then given by

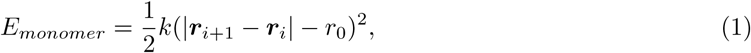

where ***r***_*i*_ is the position vector of the *i*-th monomer, and *r*_0_ = 1.2*σ* is the rest position of the spring. Note that we set *k*_*B*_*T* = 1 in the local simulation units.

All particles interact via the Lennard-Jones (LJ) potential with the other particles. The general interaction form is given by

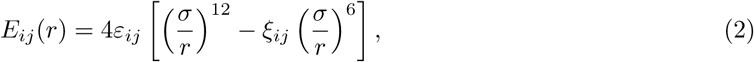

Here, the coefficients *ε*_*ij*_ and *ξ*_*ij*_ depend only on the particle identity (a, b, c or d); *r* is the center to center distance between the two particles. The parameter *ξ*_*ij*_ is set to zero for particles that interact only through hard-core repulsion, and it is set to one for particles that have an attractive component. As usual, we apply a cutoff distance for the non-bonded interactions, set to *r*_*cut*_ = 3*σ*.

The binders were assigned a fixed, strong interaction energy with the binding sites, *ε*_*bc*_ = 10*k*_*B*_*T*. The interaction between the tracers and the polymer is given by *ε* ≡ *ε*_*da*_ = *ε*_*db*_. This is the other parameter that we vary in our simulations.

We used 25 different values of *ϕ*, from 2% to 50%, in steps of 2%, and 10 different values of *ε*, from 0.9*k*_*B*_*T* to 2.7*k*_*B*_*T*, in steps of 0.2*k*_*B*_*T*. We simulated our system for all possible pairs of *ϕ* and *ε*. For each pair, ten independent simulations were carried out, each one with a different distribution of randomly placed binding sites (b sites) on the polymer.

Molecular Dynamics (MD) simulations were carried out using the HOOMD-blue software [Anderson et al., 2008, Glaser et al., 2015]. To keep the temperature of the system constant at the value *T*, the system was simulated using Langevin dynamics. To each particle of the system it was assigned a drag coefficient *γ* = 1 (in local simulation units), equal for each particle. This gives rise to a bulk diffusion coefficient *D*_0_ for free particles of *≈* 1.2cm^2^*/*s in real units (see also Appendix B).

We chose the simulation box to be cubic, with edge length *L* = 50*σ*. Periodic boundary conditions were applied in the simulations.

An equilibration of 2 10^7^ time steps was run before taking samples of the system. After that, snapshots of the system were taken every 10^4^ steps. The total length of the MD runs was 10^8^ steps.

### 4.2 Polymer contact matrix and tracer occupancy

To assess the relationship between the 3D structure of the polymer and the diffusing properties of the system, we evaluated the contact matrix of the polymer with itself. To obtain something similar to chromosome capture-based interaction matrices, we evaluated a boolean contact matrix, which gives for each pair of particles a value of 1 if the two particles are in contact, and 0 otherwise. The criterion to establish whether the two particles are in contact is by assessing whether the center to center distance between the two particles is smaller than a given threshold, *t*. Note that distances must be calculated taking into account the periodic boundary conditions of the simulation box. All the distance matrices were computed using the Python package MDAnalysis [Gowers et al., 2016, Michaud-Agrawal et al., 2011]. For each of the statistically independent snapshots of the system, we evaluated the contact matrix. The final matrix, **H**, is given by the sum of the contact matrices for each snapshot.

In a similar way to what done for the polymer contact matrix, we evaluate the contacts between the tracers (D particles) and the polymer. For each snapshot, we evaluate the contact matrix between all the tracers and all the particles that compose the polymer. We then consider as a “contact” any mutual distance shorter than *t*. The choice of *t* is somewhat arbitrary. We set this value to *t* = 2*σ*, which seems a natural choice for it is the diameter of each particle, and also a distance at which the LJ interactions are practically zero. We then can compute the tracer occupancy profile of the tracers to the polymer, which we call ***C***. This vector is obtained by summing over all the tracers of the contacts with the polymer.

### 4.3 Monte Carlo simulation of the tracers

We set up Monte Carlo (MC) simulations of particles exploring the structure of an immobile polymer. A snapshot was taken from our MD simulations at random, the tracers were removed from the snapshot, and the polymer and binder configuration were recorded.

A single particle was simulated using the Metropolis algorithm [Metropolis and Ulam, 1949]. The simulations were carried out using only the hardcore repulsion term in the Lennard-Jones potential (no *r*^−6^ term in Eq. (2), equivalent to setting *ξ*_*ij*_ = 0). The position of the particle at successive MC steps was taken randomly in the simulation box. The MC runs last 10^6^ steps.

### 4.4 Volume exclusion effects

If there are many binding sites on the polymer (high *ϕ*), its configuration in three dimensions will be compact, with the binding sites buried deep inside the globular part of the polymer. The more this is true, the more we expect the tracers to the be excluded from contacting the binding sites that are in the globular core. Instead, we expect the tracers to bind more to the outer shell of the globule, which is more enriched with non-binding sites. To test these intuitive ideas and quantitatify volume exclusion effects on the diffusion of the tracers, we use several different metrics.

First, we calculate the tracer “coverage”, defined as the percentage of monomers that are visited by the tracers during the simulation time frames. If some monomers are never contacted by the tracers because of volume exclusion, the coverage will be less than 100%.

Next, we calculate the Pearson correlation coefficient between the tracer occupancy and the binder occupancy (calculated the same way as the tracer occupancy). If the tracers are excluded away from the polymer because of the binders, we expect that the two occupancy profiles will be anticorrelated.

Finally, we defined a coefficient of enrichment of the binding of the tracers to the binding sites on the polymer as follows:

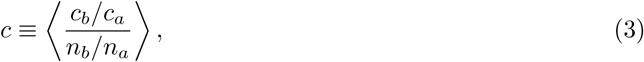

where *c*_*a*_ and *c*_*b*_ are respectively the number of binding events to a and b particles, and *n*_*a*_ and *n*_*b*_ are respectively the number of a and number of b particles on the polymer. Here, the average is performed over all the statistically independent frames of the simulations.

### 4.5 Biological data from GM12878

The Hi-C data was downloaded from GEO [Rao et al., 2014, accession ID GSE63525]. Loop positions were obtained from the file `GSE63525_GM12878_primary+replicate_Arrowhead_domainlist.txt.gz.` The locations of the highly occupied targets (HOTs) produced by Foley and Sidow [2013] were downloaded from http://mendel.stanford.edu/sidowlab/downloads/hot/analysis/. We used the file `peaks_GM12878.fasta` and obtained the positions of the domains from the headers. The overlap between the two was computed with custom R scripts [R Development Core Team, 2008] using the package GenomicRanges [Lawrence et al., 2013].

To compute the rates of contact decay, the raw Hi-C reads were normalized in 50 kb windows using the TADbit pipeline [Serra et al., in press] with default parameters. The contact decay were estimated using a simple linear regression between the logarithm of the linear separation between the windows and the logarithm of the normalized Hi-C signal. For each window, only the closest 20 windows on each side were used for the regression, as the signal in windows beyond the twentieth was typically too noisy. Also, the self contacts in the window were discarded for being overly influential on the regression parameters. The slope of the regression line served as estimate for the rate of contact decay.

## Acknowledgements

R.C is grateful to Jean-Marc Victor, the members of the CNRS GDR 3536 (ADN), and Francesco Alessandro Massucci for useful discussions. The authors would like to thank Egor Tiavlovsky for his help on an early version of the manuscript and the CRG Scientific Information Technologies for helping with the simulation setup.

This work was supported by the Spanish Ministry of Economy and Competitiveness, Centro de Excelencia Severo Ochoa 2013-2017, SEV-2012-0208, Plan Nacional BFU2012-37168 ERC Synergy Grant 609989, and the People Programme (Marie Curie Actions) of the European Union’s Seventh Framework Programme (FP7/2007-2013) under REA grant agreement n 608959. We acknowledge the support of the CERCA Programme / Generalitat de Catalunya.

## Appendix A Polymer conformation

We investigated the properties of the polymer configuration in our simulation sets. First, we looked at the radius of gyration *r*_*gyr*_ of the polymer, which is defined as follows:

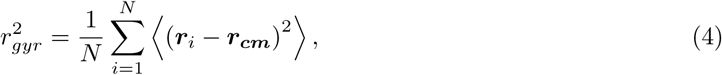

where ***r*_*cm*_** is the center of mass of the polymer. Figure 8A-B shows the average values of the radius of gyration, calculated by averaging over the last 5 *·* 10^7^ trajectory frames of the simulations. In the case of *n*_*t*_ = 10, the radius of gyration remains roughly constant as *ε* increases. However, when *n*_*t*_ = 200 there is a sharp decrease of the radius of gyration when increasing *ε*. This is a sign that the tracers are driving a coil-globule transition in the polymer.

**Figure 8:**
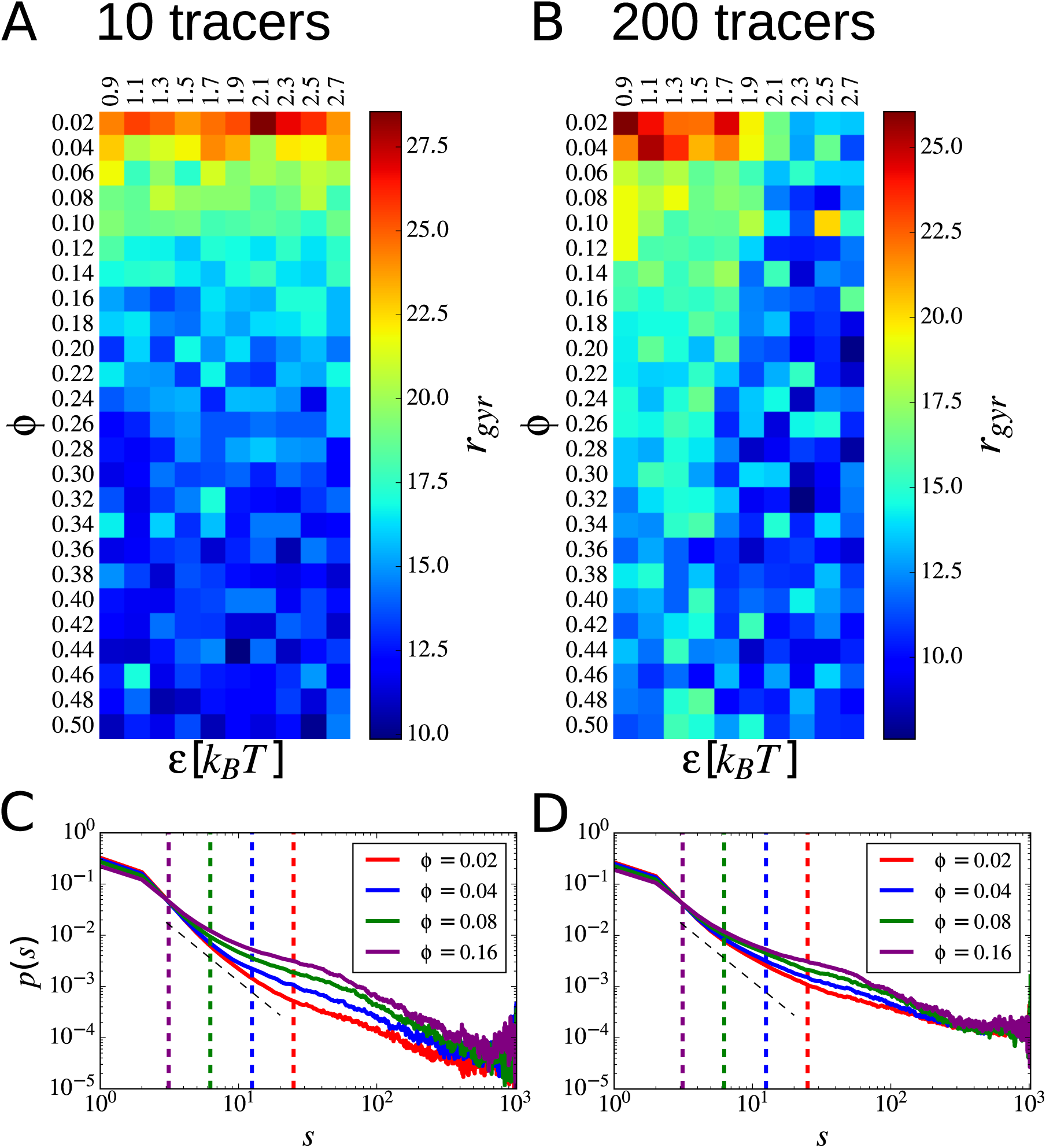
(A-B) Average radius of gyration as a function of the two parameters *phi* and *α*. (C-D) Average probability of contact *p*(*s*) as a function of the linear distance along the polymer *s*. Curves are shown for four different values of *ϕ*. The vertical dashed lines correspond to the length of the shortest half-loop *l* at the given value of *ϕ* (see main text).

Next, we examined the probability of contact *p*(*s*), which represents the average probability that any given monomer is in contact with any other which is *s* monomers away. From classical polymer physics arguments, we expect that *p*(*s*) ∼ *s*^*α*^, where *α* is an exponent that depends on the polymer interactions and on the particular state in which the polymer is found. Figure 8C-D shows the average values of *p*(*s*) depicted for a few different values of *ϕ* and *n*_*t*_ = 10, 200. For a given value of *ϕ*, we can define the average minimal loop size *l* as the average distance between binding sites, that is *l* = *Nϕ*. We expect then that up to *s* ≈ *l* the *p*(*s*) function will be dominated by the contribution due to this minimal loop. It turns out [Uehara and Deguchi, 2013] that up to half of this distance the exponent *α* ≈-2.20. There is a good agreement between the expected decay and the one observed in our simulations.

## Appendix B Diffusion of the tracers

To assess the diffusive properties of the tracers, we calculated the mean square displacement (MSD) of the tracers, as follows:

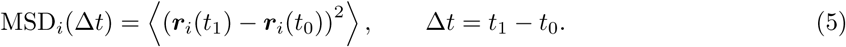

Here ***r***_*i*_ is the position vector of the *i*-th tracer, and the average ⟨*…* ⟩ is performed over the statistically independent snapshots. The calculation of the MSD was performed by averaging over many possible independent starting times *t*_0_, following the guidelines in Ref. [Qian et al., 1991].

For sufficiently long time intervals, the values of the MSD as a function of time obey the relationship

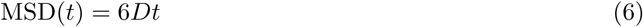

where *D* is the diffusion coefficient. For a tracer that does not have any interactions with other particles in the system, we expect that the value of the diffusion coefficient should be given by the Einstein relationship:

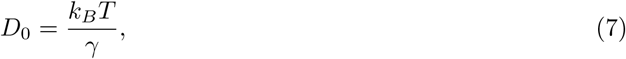

where *γ* is the drag coefficient of the particle.

In general however it is interesting to look at the *instantaneous* diffusion coefficient, which is obtained from the MSD time trace by taking its time derivative:

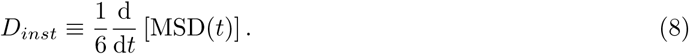

For each of the diffusing tracers we expect a different value of the instantaneous diffusion coefficient, because each tracer will randomly bind and unbind to the polymer in the simulation. Therefore, the most useful representation of the average properties of the diffusion in each simulation is the plot of the distribution of instantaneous diffusion coefficients, as the parameters in the simulations vary.

We investigate the relationship between the diffusion properties of the tracers and their affinity to the polymer, together with the three-dimensional structure of the polymer. Figure 9 summarizes our finding on the diffusion behavior of the tracers. Figure 9A shows an the MSD curves as a function of time for the ten tracers in an example simulation. For the chosen value of *ε* = 2.1*k*_*B*_*T*, the tracers have markedly different behaviors: some of them have a steep MSD curve with slope close to the value of the theoretically expected diffusion coefficient *D*_0_ (see Eq. (7)), whereas other tracers are almost immobile (flat MSD(*t*) curve).

**Figure 9:**
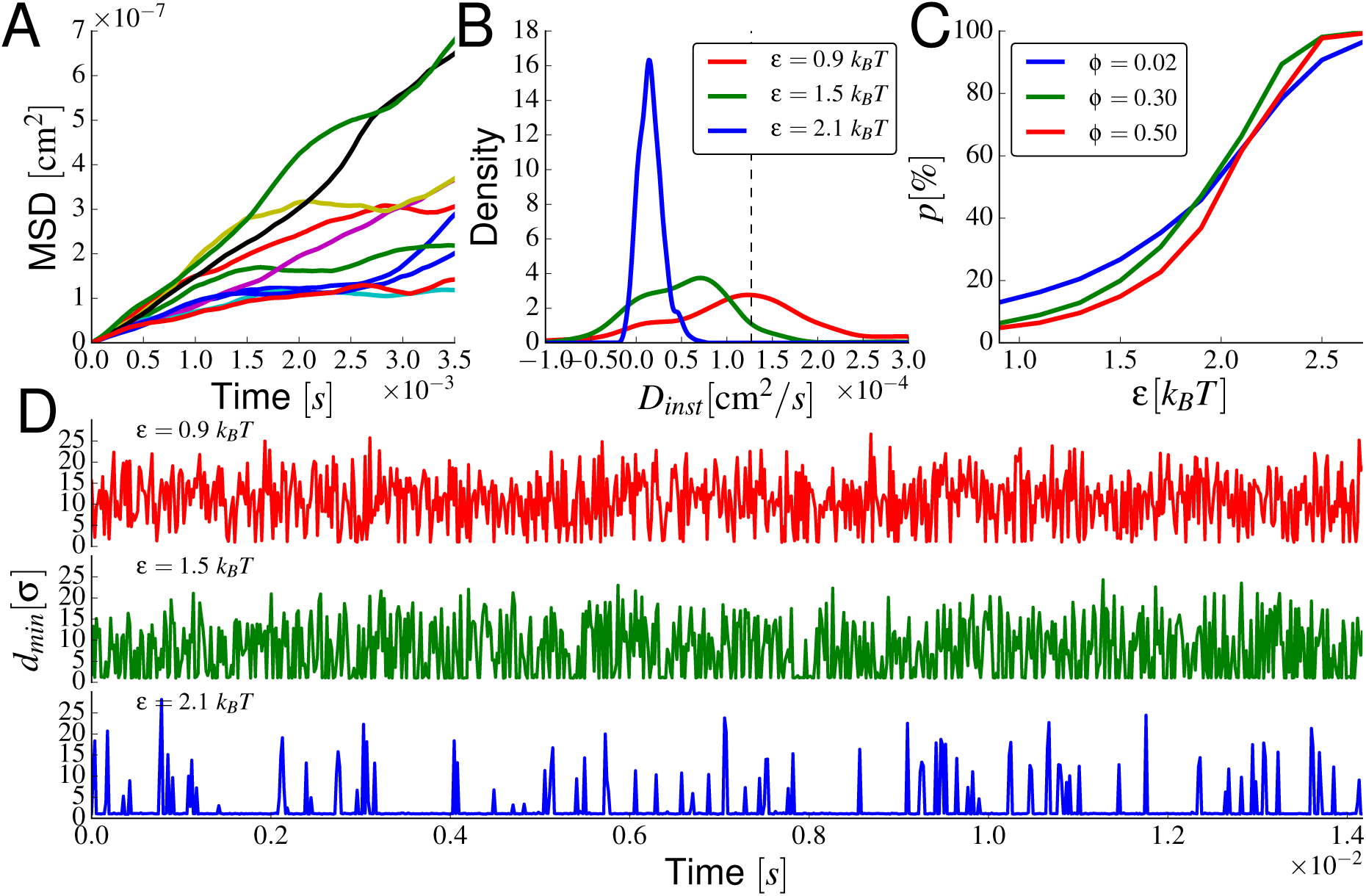
Diffusion properties of the tracers in the simulations. (A) MSD as a function of time for the tracers in an example simulation at *ϕ* = 0.2, *ε* = 2.1*k*_*B*_*T* and *n*_*t*_ = 10. (B) Distribution of instantaneous diffusion coefficients for three example simulations at different values of *ε* = 0.9, 1.5, 2.1*k*_*B*_*T* and *ϕ* = 0.20. Dashed vertical line corresponds to the value of *D*_0_ (see Eq. (7)). (C) Percentage of binding events *p* as a function of the tracer-polymer affinity, for three different values of *ϕ*. The value of *p* corresponds to the average fraction of simulation frames in which the minimum distance between the tracer and the polymer *d*_*min*_ is lower than the Lennard-Jones cutoff length *r*_*cut*_. (D) Three example time traces of *d*_*min*_ as a function of the simulation time, for three different values of the tracer-polymer affinity *ε*.

**Figure 10:**
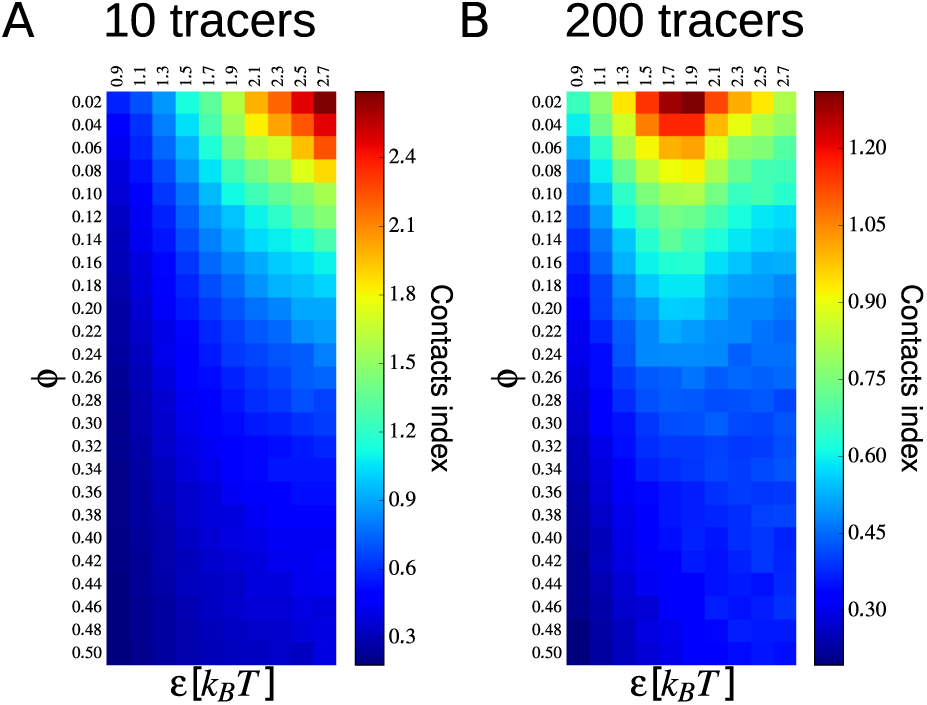
The average contacts enrichment coefficient *c* (defined in Eq.(3)), plotted for *n*_*t*_ = 10 (A) and *n*_*t*_ = 200 (B).

To characterize further this behavior, we calculated the distribution of instantaneous diffusion coefficients. Figure 9B shows the distribution of *D*_*inst*_ for three different simulations, for a fixed value of *ϕ* = 0.20. As *ε* grows, the instantanous diffusion coefficients become smaller. There is however always a fraction of the tracers that has instantanous diffusion coefficient close to *D*_0_ (dashed vertical line in the figure).

We also investigated the effect of the three-dimensional structure of the polymer on the diffusion properties of the tracers. To this end, we calculated the minimum distance between a tracer and the polymer (*d*_*min*_), and the percentage *p* of frames in the simulations in which *d*_*min*_ is smaller than *t*, the threshold for the definition of a contact. Figure 9C shows the variation of *p* as a function of *ε*, for three fixed values of *ϕ*. In every case, there is a monotonic increase of the percentage of binding events when *ε* increases. The same can be seen in figure 9D, where the time traces of three example tracers are plotted, for three different values of *ε*. The effect of the three-dimensional structure of the polymer, as proxied by the value of *ϕ*, is less pronounced than the effect of *ε*. Figure 9C shows that lower values of *ϕ* correspond to higher values of *p*, but only for small tracer-polymer affinity. This is again an indication that high values of *ϕ* hinder binding of the tracers.

